# Sensitive detection and propagation of brain-derived tau assemblies in HEK293 based wild-type tau seeding assays

**DOI:** 10.1101/2024.07.18.604077

**Authors:** Melissa Huang, William A McEwan

**Affiliations:** UK Dementia Research Institute at the University of Cambridge, Department of Clinical Neurosciences, Hills Road, Cambridge, CB2 0AH, United Kingdom

## Abstract

The assembly of tau into filaments defines the tauopathies, a group of neurodegenerative diseases including Alzheimer’s disease (AD), Pick’s disease (PiD), corticobasal degeneration (CBD) and progressive supranuclear palsy (PSP). The seeded aggregation of tau has been modelled in cell culture using pro-aggregant modifications such as truncation of the N- and C-termini and point-mutations within the tau microtubule-binding repeat domain. While providing experimental convenience, this limits the applicability of research findings to sporadic disease, where filaments contain wild-type, full-length tau isoforms. We therefore aimed to develop a sensitive and specific biosensor assay for the seeded aggregation of brain-derived tau species utilizing full-length, wild-type tau stably expressed in HEK293 cells. We show that addition of brain-derived tau extracted from cases of AD, PiD, CBD, or PSP induces the formation of HA- or GFP-tagged 0N3R or 0N4R tau aggregates. By isolating and expanding a single cell containing AD-seeded aggregates, we generated a cell line that propagates insoluble tau. We demonstrate that HEK293-propagated tau is hyperphosphorylated at disease relevant sites and retains a seeding profile similar to AD brain-derived material. We propose that these cell lines will aid pre-clinical, high-throughput screening for modifiers of seeded aggregation with greater conformational and strain specificity than existing cell-based biosensor assays.

## Introduction

Microtubule-associated protein tau assembles into highly ordered amyloid filaments in a subset of neurodegenerative diseases, termed tauopathies. Mutations in the autosomal dominant *MAPT* gene are causative of tau aggregation and subsequent neurodegeneration in frontotemporal dementia and parkinsonism linked to chromosome 17 (FTDP-17) (Hutton et al., 1998; Poorkaj et al., 1998; Spillantini et al., 1998). Studies utilizing these *MAPT* mutations have firmly established the importance of abnormal tau assembly to neuronal dysfunction and death. Mouse models transgenic for human P301S tau, a mutation causative of early-onset FTDP-17 (Bugiani et al., 1999; Lossos et al., 2003; Sperfeld et al., 1999; Werber et al., 2003; Yasuda et al., 2005; Yasuda et al., 2000), develop hyperphosphorylated, filamentous tau inclusions and neurodegeneration (Allen et al., 2002; Macdonald et al., 2019; Yoshiyama et al., 2007). Importantly, the majority of neurodegenerative tauopathies develop independently of *MAPT* mutations. For example, accumulation of hyperphosphorylated tau aggregates occurs in Alzheimer’s disease (AD), sporadic Pick’s disease (PiD), corticobasal degeneration (CBD), and progressive supranuclear palsy (PSP) in absence of *MAPT* mutations. The accumulation of tau aggregates in cases of “wild-type” tauopathies have nevertheless been shown to correlate with brain atrophy and the severity of dementia (Arriagada et al., 1992; La Joie et al., 2020).

Six isoforms of tau are expressed by alternative splicing of the *MAPT* gene in the central nervous system of healthy adults. Three isoforms exclude exon 10, which encodes the second repeat (R2) of the microtubule binding repeat domain (3R tau), and three isoforms include all four repeat domains (4R tau) (Goedert et al., 1989). Recent electron cryo-microscopy structures have shown that the core of the amyloid filaments found in tauopathies is comprised of the microtubule binding repeat domains, while the N-and C-termini of tau protein remains largely unstructured (Scheres et al., 2023). Consequently, the isoform composition of tau filaments can differ between different diseases. In AD, a mixture of 3R and 4R tau isoforms are present in the filaments, whereas 3R tau makes up the filaments found in PiD and 4R tau is found in CBD and PSP.

Studies of recombinant, purified tau protein have demonstrated that the wild-type full-length protein is highly soluble, and requires the addition of negatively charged polyanionic co-factors (Friedhoff et al., 1998; Goedert et al., 1996; Kampers et al., 1996) or agitation with glass beads (Chakraborty et al., 2021) to form filaments *in vitro*. Tau filaments induced to form with heparin (Zhang et al., 2019), phosphoserine, or RNA (Lovestam et al., 2022), have been shown to form structures distinct from those found in human disease. Recent advances have shown that, under specific assembly conditions, tau fragments consisting of the microtubule binding repeats can be induced to form paired helical filaments (PHFs) structurally identical to those found in AD (Lovestam et al., 2022). Full-length tau assemblies, however, make up the aggregates found in tauopathies but, to date, the spontaneous formation of full-length *in vitro* assembled PHFs has not yet been reported.

The seeded aggregation of full-length tau has been shown experimentally in various cell culture models, in which brain-derived or recombinant tau assemblies can enter the cellular environment to seed the subsequent aggregation of native tau (Frost et al., 2009; Guo & Lee, 2011; Kfoury et al., 2012). Such experiments have further shown that the conformation of the tau seeds dictates the potency of seeded aggregation (Falcon et al., 2015), and that tau isolated from a 3R or 4R tauopathy specifically seeds mutant repeat domains of 3R or 4R tau, respectively (Woerman et al., 2016). Further, mutant tau assemblies can preferentially promote aggregation of monomer bearing the same mutation over wildtype (Aoyagi et al., 2007). Despite this, the most commonly used “biosensor” assays for seeding competency of tau species rely on the P301S mutation on either truncated (Holmes et al., 2014) or full-length (McEwan et al., 2017) tau. The conformations resulting from seeding onto the P301S tau expressed in cell culture is unknown, however, given the recent finding that transgenic mouse models expressing human P301S tau develop tau filament structures distinct from that found in human disease (Schweighauser et al., 2023), it is likely that the structure of the original seed is not accurately propagated. On the other hand, Tarutani et al. demonstrated that full-length, wild-type 1N3R or 4R tau transiently expressed in undifferentiated SHSY5Y neuroblastoma cells could be seeded by brain-derived seed to propagate tau filament structures that maintain some structural properties of the original filaments found in AD and CBD.

We here develop a biosensor assay using HEK293 cells overexpressing full-length, wild-type 0N3R or 0N4R tau, which responds sensitively to brain-derived tau seeds and exhibits isoform-specificity for either 3R or 4R tauopathies. Further, we describe a cell line which retains hyperphosphorylated, AD-propagated insoluble tau over multiple passages. We demonstrate the use of these aggregate-containing HEK293 cells as a source of seed-competent tau assemblies which harbour PTMs similar to those found in disease. We show that our cell lines respond preferentially to AD brain-derived or AD HEK-propagated tau seeds over *in vitro* heparin-induced filaments, suggesting that the HEK-derived tau retains biological properties of AD brain-derived tau.

## Results

We developed a HEK293 cell line stably expressing the 0N3R or 0N4R isoform of human tau with an N-terminal HA (YPYDVPDYA) tag by lentiviral transduction. Sarkosyl-insoluble tau assemblies extracted from AD patient brain (AD seed) were introduced to the cells using Lipofectamine 2000. Addition of AD seed resulted in the formation of methanol-resistant HA-tau puncta which co-localize with an antibody selective for hyperphosphorylated, pathological tau (AT100), by immunofluorescence microscopy (Figure 1A). We observed a seed concentration dependent increase in the percent of cells developing HA-tau puncta, with no puncta observed in the untreated control cells. Significant seeding of HA-tau puncta was detectable in the HA-0N3R expressing cell line at 0.16 ng/mL of AD seed, which is equivalent to approximately 0.2 μg of brain, extracted, per well. We observed a significant difference in the seeding competency of HA-0N3R compared to HA-0N4R tau in response to AD seed at all concentrations where seeding was detected (p < 0.0001), consistent with previous reports in transiently transfected, undifferentiated SHSY5Y cells (Tarutani et al., 2023; Tarutani et al., 2021) (Figure 1B). This was despite the HA-0N4R expressing slightly more total tau than the HA-0N3R cell line (p = 0.0278) (Figure 1C). We confirmed that the appearance of immunoreactive puncta corresponded to the accumulation of sarkosyl-insoluble HA-tagged tau upon the addition of seed (Figure 1D).

**Figure 1.**
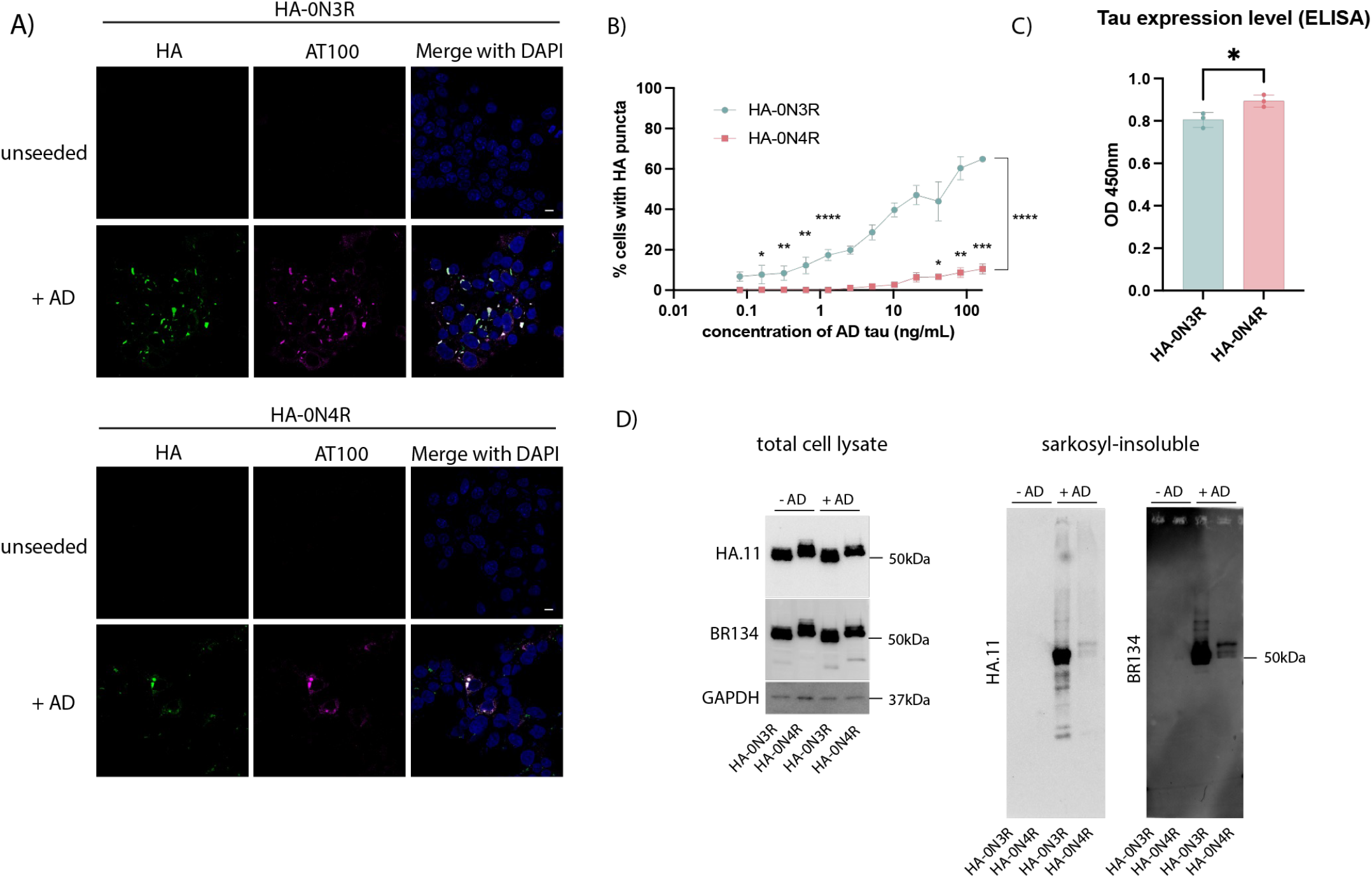
Seeded aggregation of HA-tagged full-length, 0N3R and 0N4R wild-type tau by AD brain-derived tau seed. A) Representative immunofluorescence images of HEK293 cells stably expressing HA-0N3R or HA-0N4R tau with or without treatment AD seed in the presence of Lipofectamine 2000. sb = 10μm B) Quantification of percent cells bearing HA-puncta (two-way ANOVA with Fisher’s LSD, **** p<0.0001, *** p = 0.0002, ** p <0.006, * p < 0.03). C) Total tau (HT7) ELISA showing relative expression levels of HA-0N3R and HA-0N4R HEK293 cell lines (unpaired t-test, * p = 0.0278). D) Immunoblot of total cell lysate and sarkosyl-insoluble fraction of HEK293 cell lines with or without treatment with AD seed using a C-terminal tau antibody (BR134) and anti-HA antibody.

Clonal populations of cells were isolated from HA-0N3R expressing cells seeded with AD brain by limiting dilution. Clones were screened by immunofluorescence microscopy for persistent HA-tagged puncta, and a positive clone was selected and expanded (HA-0N3R^AD^). HA-0N3R^AD^ HEK293 cells contain HA-tagged tau aggregates which are hyperphosphorylated at pT231 (AT180), pS202/pT205 (AT8), pT212/pS214 (AT100), pS396, pS422, and are positive for AmyTracker 680, an amyloid-specific dye (Figure 2A). These cells maintain insoluble tau aggregates over multiple passages, though they exhibit a decrease in the amount of insoluble tau over extended passage (Figure 2B,C). To investigate the utility of these cells as a source of hyperphosphorylated tau seeds, we incubated eGFP-0N3R or 0N4R expressing HEK293 cells with sarkosyl-insoluble tau extracted from AD brain or HA-0N3R^AD^ HEK293 cells (HA-0N3R^AD^ seed). The use of a distinct tag (eGFP) ensures that the original HA-0N3R^AD^ seed is not the source of signal. In both cell lines, AD seed and HA-0N3R^AD^ seed behaved similarly, with equal amounts of assembled tau inducing similar percentages of seeded cells, as measured by eGFP puncta, at all concentrations (p > 0.05) (Figure 3A). Heparin-assembled tau filaments are frequently used a source of tau seeds in place of brain-derived material to study the seeded aggregation of tau. While heparin-assembled recombinant 0N3R, 0N4R, and 0N4R-P301S tau (0N3R^heparin^, 0N4R^heparin^, 0N4R-P301S^heparin^) seeded aggregation in cells expressing eGFP-0N4R-P301S tau, we observed that only 0N3R^heparin^ was able to seed aggregation in eGFP-0N3R cells. Both 0N3R^heparin^ and 0N4R^heparin^ tau assemblies were able to seed aggregation in eGFP-0N4R cells, albeit requiring higher concentrations than for AD seed or HA-0N3R^AD^ seed. 0N4R-P301S^heparin^ assemblies were incompatible with seeding in both eGFP-0N3R and 0N4R cells. HA-0N3R^AD^ seed exhibited a greater level of seeding than heparin-assembled tau at all concentrations where seeding was detected (Figure 3A). All three cell lines expressed similar levels of total tau (Figure 3B).

**Figure 2.**
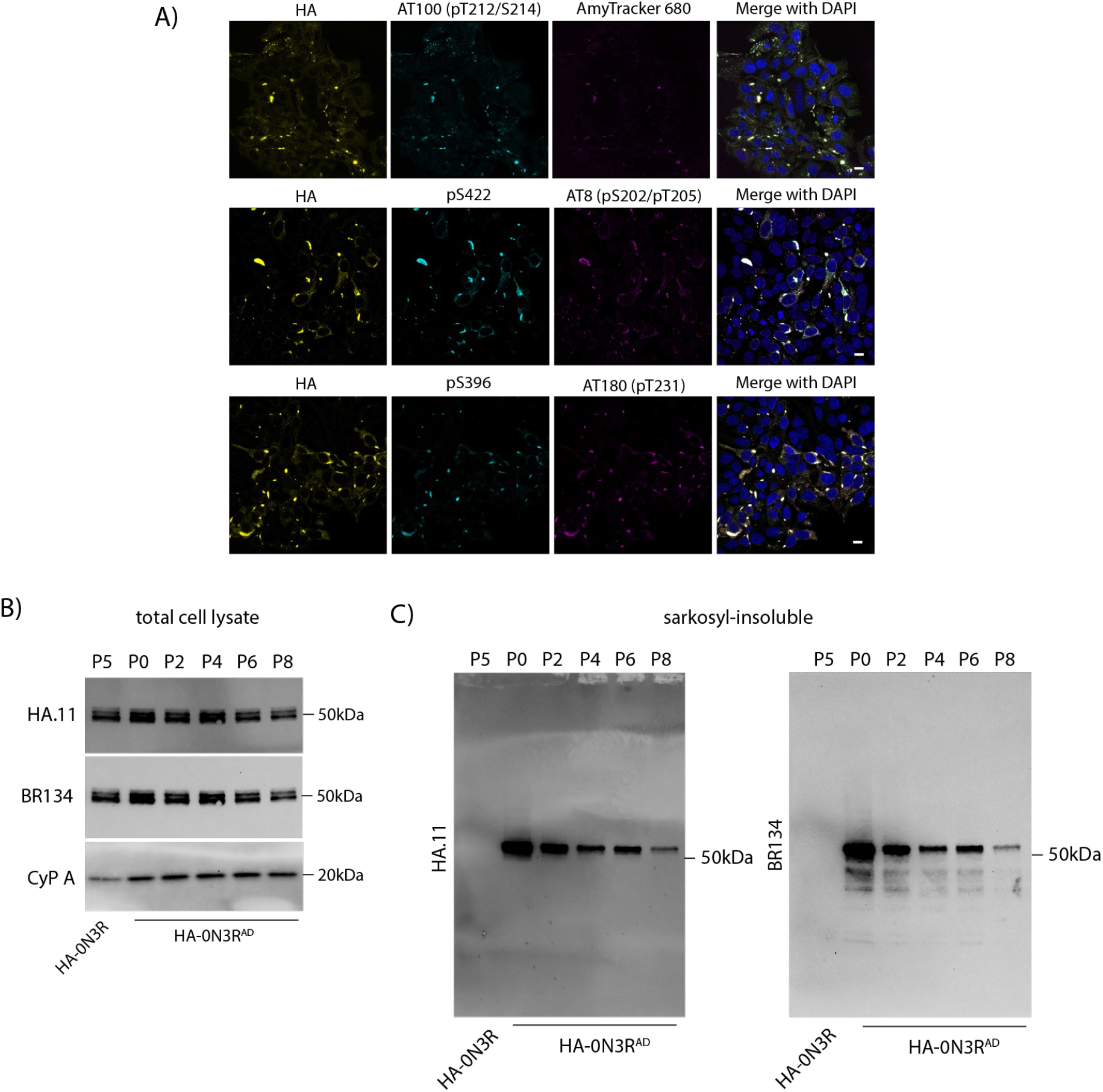
HA-0N3R^AD^ HEK293 cells propagate AD-brain derived tau seeds. A) Representative immunofluorescent images of HA-0N3R^AD^ cells at passage 4 (P4), probing with phosphorylation-specific antibodies: AT180, AT8, AT100, pS396, and pS422, and with an amyloid specific dye, Amytracker 680. sb = 10μm B). Immunoblot of total cell lysate of HA-0N3R at passage five (P5) and HA-0N3R^AD^ cells over multiple passages. C). Immunoblot of the sarkosyl-insoluble fraction of HA-0N3R (P5) and HA-0N3R^AD^ cells over multiple passages using a C-terminal tau antibody (BR134) and anti-HA antibody.

**Figure 3.**
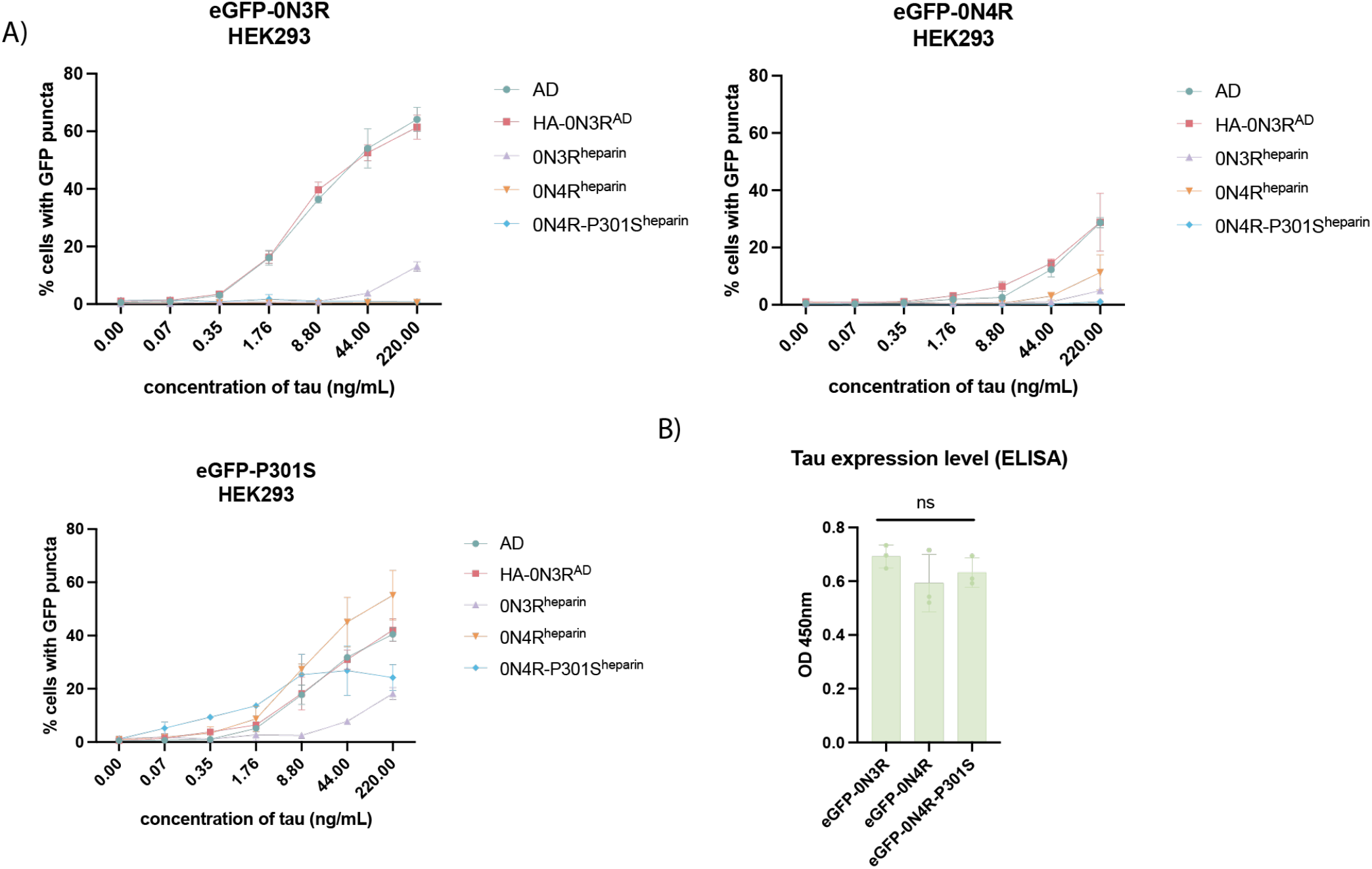
WT-tau expressing cell lines respond preferentially to AD-brain derived seed A) Levels of seeded aggregation in HEK293 cells expressing the indicated eGFP-tagged tau construct following treatment with AD-brain derived tau assemblies, with AD-HEK derived tau (HA-0N3R^AD^) or with recombinant heparin tau assemblies (0N3R^heparin^, 0N4R^heparin^, 0N4R-P301S^heparin^). B) Total tau ELISA showing relative expression levels of eGFP-0N3R, eGFP-0N4R, and eGFP-0N4R-P301S HEK293 cells (One-way ANOVA with Tukey’s multiple comparisons test, p > 0.05).

While the AD filament fold excludes the R1 and R2 domain of the microtubule binding repeat domains, and can thus incorporate both 3R and 4R isoforms in its structure, other sporadic tauopathies have filament structures which allow incorporation of only 3R (PiD) or 4R (CBD, PSP) tau isoforms (Shi et al., 2021). Isoform-specific seeded tau aggregation of full-length, wild-type tau has been shown previously in transiently transfected SH-SY5Y cells (Tarutani et al., 2023). We aimed to show that our HEK293 stable cell lines also respond with isoform specificity to 3R or 4R tau strains. We extracted sarkosyl-insoluble tau from AD, PiD, CBD, and PSP patient brain tissue, and confirmed the presence of 3R or 4R insoluble tau by immunoblotting (Figure 4A). Addition of sarkosyl-insoluble tau extracted from PiD patient brain specifically seeded the eGFP-0N3R cell line (Figure 4B) and tau extracted from CBD or PSP patient brain specifically seeded the eGFP-0N4R cell line (Figure 4B). We confirmed that the cells develop sarkosyl-insoluble GFP-tagged tau upon the addition of seed (Figure 4C).

**Figure 4.**
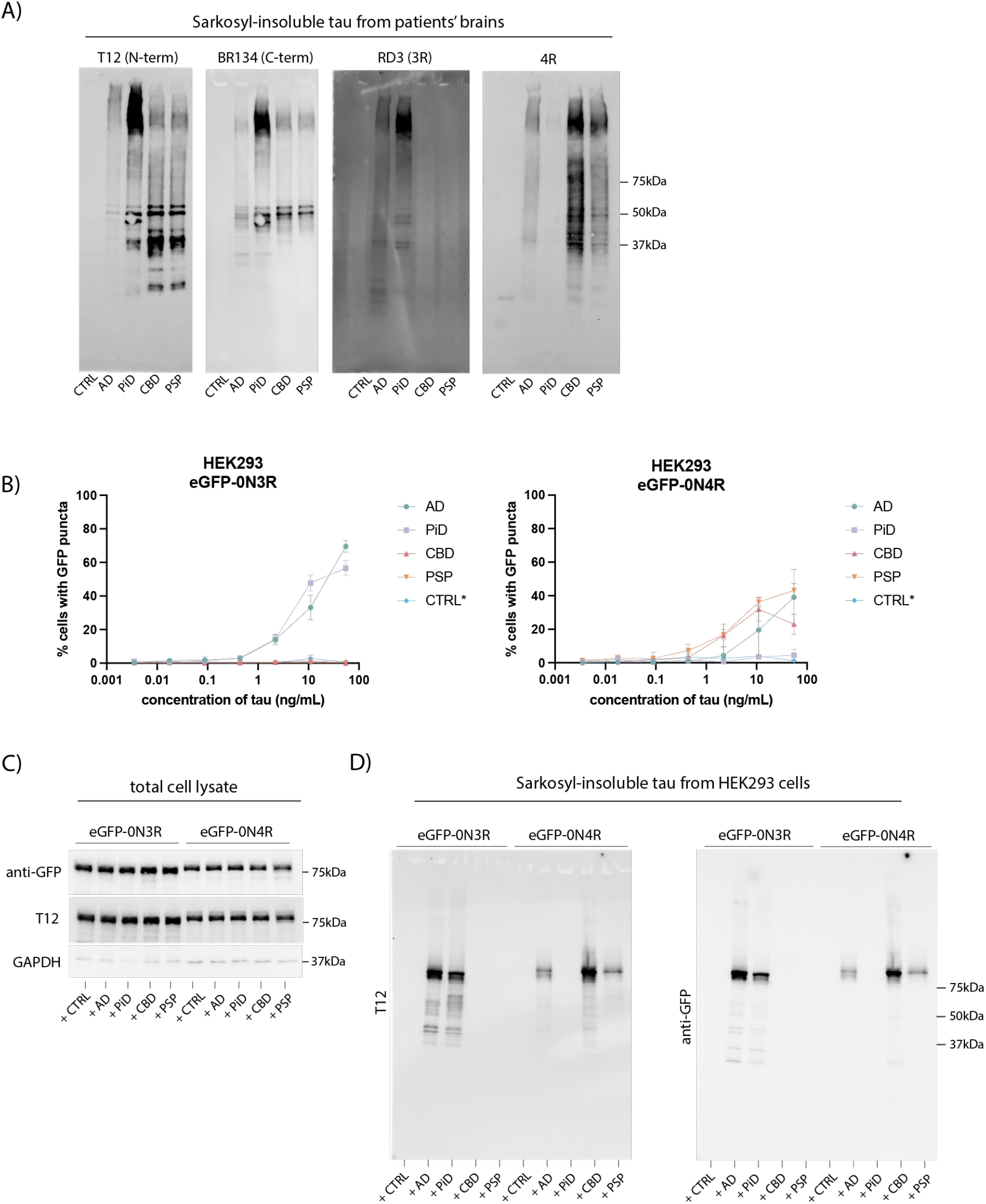
Wild-type tau expressing cell lines exhibit isoform specificity for 3R or 4R tauopathies. A) Immunoblotting of the sarkosyl insoluble fraction from control, AD, PiD, CBD, and PSP brain tissue using antibodies against the N-terminus (T12) and C-terminus (BR134) of tau, 3R tau (RD3) and 4R tau (anti-4R). B). HEK293 cells engineered to express either 0N3R or 0N4R tau with an N-terminal eGFP tag were treated with sarkosyl-insoluble fractions prepared from control (CTRL), AD, PiD, CBD, and PSP brains in the presence of Lipofectamine 2000. Concentration of brain-origin insoluble tau was normalized following quantification using the total tau ELISA kit (ab273617). * For the control brain extract, the mean volume of seed for the disease brain extracts was used, as no sarkosyl-insoluble tau was detected by ELISA. C). Immunoblot analysis of total cell lysates and D) sarkosyl-insoluble fractions of eGFP-0N3R and eGFP-0N4R HEK293 cells treated with control, AD, PiD, CBD, and PSP brain extract using antibodies against the N-terminus (T12) of tau and anti-GFP.

## Discussion

The diversity of possible tau conformations has been revealed to be large: in addition to the eight known wild-type tau structures associated with human disease (Scheres et al., 2023) around fifty conformations of tau, or its truncation products, have been reported to arise during in vitro aggregation (Lovestam et al., 2022; Lovestam et al., 2024; Zhang et al., 2019). The biological significance of these structures remains unclear, but the observation that particular folds are associated with specific diseases raises the possibility that tau filament conformation is closely tied to its pathogenesis. Given that mutations in the amyloid core region of tau are likely to have a consequential effect on the structure of tau filaments (Schweighauser et al., 2023), methods that enable propagation of wild-type tau are required to effectively model brain-derived tau species. Here we have developed a biosensor assay in the widely used HEK293 cell line, where wild-type tau responds to exogenously supplied tau assemblies extracted from post-mortem patient brain tissue. Aggregation is highly sensitive, with tau isolated from 0.2 μg of AD brain tissue sufficient to yield significant seeding. This finding is surprising, given the widely held view that tau mutants or truncations are required to observe seeded aggregation, particularly at low concentrations of supplied tau seeds. In contrast to cell lines expressing mutant tau, our wild-type tau expressing reporters lines respond preferentially to brain-derived tau assemblies over heparin-assembled recombinant tau. This suggests that our wild-type cell lines have greater specificity for disease-relevant conformations, while P301S cell lines are less specific and thus less likely to retain structural information from the initial seed.

Full-length tau is highly soluble, and the assembly of recombinant tau into seed-competent filaments for experimental procedures is most commonly achieved using anionic cofactors such as heparin or arachidonic acid (Goedert et al., 1996; King et al., 2000; Perez et al., 1996). The truncated core of the AD protofilament (amino acids 297-391) has been shown to form paired helical filaments (PHFs) identical to those found in AD brain (Al-Hilaly et al., 2020; Lovestam et al., 2022), but, to date, full-length recombinant tau has not been reported to form disease-relevant structures.. A possible explanation for this is that recombinant tau assemblies suffer from a lack of post-translational modifications, which are highly enriched in patient brains (Wesseling et al., 2020) and likely play an important role in tau aggregation. Generating a brain-independent source of post-translationally modified tau seeds is thus an important step to modelling and understanding tau aggregation. Unlike *Escherichia coli*, HEK293 cells can produce protein with extensive post-translational modification. Previous work has demonstrated that aggregated tau may be inherited from mother to daughter cell in HEK293 cells expressing tau residues 244-372 (Sanders et al., 2014). We here extend these findings by demonstrated that wild-type HA-0N3R tau can propagate hyperphosphorylated, insoluble tau assemblies during subsequent rounds of cell division following addition of brain-derived tau assemblies. Importantly, the extracted tau from these cells behaves near-identically to AD brain-derived tau in its ability to further induce seeded aggregation. Using phosphorylation-dependent antibodies, we found that aggregates in these cells also retain a phosphorylation status consistent with insoluble tau found in AD patient brain (Wesseling et al., 2020). Our study is the first demonstration that full-length, wild-type tau can be propagated in this way. While our findings are consistent with structural properties of AD tau being retained, one limitation is that their structure is currently unknown. Tarutani et al. (2023) showed that undifferentiated SH-SY5Y cells, transiently transfected to express full-length 1N3R and 1N4R, and seeded with AD formed a single protofilament of the AD fold. We hypothesize that the structure propagated in HEK293 cells will be similar, however further structural studies should be performed.

In conclusion, we demonstrate that full-length, wild-type tau expressed in HEK293 cells can be induced to aggregate upon the addition of brain-derived tau seeds. This system offers an alternative to P301S-based assays, which respond with less specificity to disease-relevant tau conformers. These reporter cell lines will more faithfully model wild-type tauopathies and can be used in pre-clinical settings to screen for modifiers of seeded aggregation. These cell lines can additionally be used to sensitively quantify the seeding-competency of tau species derived from human brain in different anatomical regions and at different stages of disease. We further demonstrate that HA-0N3R^AD^ HEK293 cells can be used as a source of brain-propagated seed which is hyperphosphorylated and seed-competent. Unlike recombinant assembled tau, these aggregates are hyperphosphorylated and are bound by phosphorylation-specific antibodies, which will be valuable for molecular diagnostics and therapeutics research targeting phosphorylated tau.

## Methods

### Cloning

Cloning was performed using In-Fusion (Takara) or restriction cloning. Constructs encoding 0N3R or 0N4R tau were tagged at the N-terminus with HA or eGFP tag. The pSMPP backbone (Addgene #104970) was modified by replacing the puromycin resistance gene for a neomycin resistance gene (pSMPN). Tagged-tau was subcloned into the lentiviral backbone by digestion at Mlu1/Not1 restriction sites.

### Human embryonic kidney cell maintenance

HEK293 and HEK293T cell lines were maintained in Gibco DMEM + Glutamax + 10% FBS + 1% penicillin-streptomycin in a tissue culture incubator at 37°C with 5% CO_2_

### Lentivirus production

Lentivirus was produced in HEK293T cells mediated by Lipofectamine 3000 (Invitrogen, L3000001) according to the manufacturer’s instructions for lentivirus production using pMD2.g envelope (Addgene #12259), pCRV-gag-pol packaging plasmid (a gift from Prof Stuart Neil, Kings College London) and pSMPP or pSMPN transfer plasmid backbone. HEK293 cells were transduced for 72 hours mediated by polybrene (Merck, TR10003) before selection using 1ug/mL puromycin dihydrochloride (Gibco, A1113803) or 0.25mg/mL geneticin (Gibco, 10131027). Cell lines were made clonal by limiting dilution.

### Preparation of sarkosyl-insoluble tau from human brain samples

Postmortem brain tissue and patients were obtained from the Cambridge Brain Bank under the ethically approved protocol: REC 16/WA/0240 (PSP and CBD) and NRES/20/EE/0183 (PiD). The donors were a 37 year old male dying of renal failure secondary to type 1 diabetes, with a neuropathological diagnosis of “normal”, a 74 year old female with confirmed neurobiological diagnosis of CBD, and an 85 year old male with confirmed neuropathological diagnosis of PSP. Postmortem brain from a patient with AD was obtained from the Oxford Brain Bank (Ethics approval reference: 15/SC/0639), UK Sound Central Oxford C Research Ethics Committee. The donor was an 81 year old female with AD neuropathological diagnosis. Frozen brain tissue from frontal cortex (1.2-2.4g) was homogenized in 10X (v/w) of extraction buffer containing 800mM NaCl, 10mM Tris-HCl pH 7.4, 2.5mM EDTA, 10% sucrose, 2% sarkosyl + Pierce Protease and Phosphatase Inhibitor Mini Tablets (Thermo Scientific, A32959), using T 10 basic ULTRA-TURRAX, (IKA 9993737002). Tissue homogenate was incubated at RT for 1 hour on a roller, then centrifuged at 13,000g for 20 mins at RT (room-temperature). Supernatant was filtered through a 0.45μm cell strainer and transferred to 1.5mL ultracentrifuge tubes, then spun at 124,500g for 2 hours at RT in a TLA-55 rotor and Beckman Coulter Optima MAX-XP ultracentrifuge. Pellets were combined and resuspended in 1mL extraction buffer and re-centrifuged at 124,500g for 1 hour. Pellet(s) was [combined and] resuspended in 1mL 50mM Tris-HCl, 150mM NaCl. Resuspended pellet was re-ultracentrifuged as before, and resulting pellet was resuspended in 250uL/g starting weight and sonicated in a water-bath sonicator for 20 cycles of 15 sec on/ 5 sec off. Concentration of sarkosyl-insoluble tau was determined by Human Tau ELISA Kit (ab273617).

### Preparation of sarkosyl-insoluble tau from HEK293

Media was removed and cells were washed 2x with PBS. Cells were lysed in 250μL / well of a 6-well plate of cell extraction buffer (800mM NaCl, 10mM Tris-HCl pH 7.4, 2.5mM EDTA, 15% sucrose, 1% sarkosyl + Protease and Phosphatase Inhibitor (Pierce)) and collected by repeated pipetting. Cells were further lysed by sonication with a metal probe Q55 sonicator (30% amplitude, 15 seconds), before incubating at 37°C for 30 minutes with shaking. Lysate was spun at 124,500g for 1 hour at RT in a TLA-55 rotor. Pellet was washed 2x in TBS before resuspending in the final volume (6uL of TBS per well of a 6-well plate). Concentration of sarkosyl-insoluble tau was determined by Human Tau ELISA Kit (ab273617).

### Immunoblotting

Cell lysate or sarkosyl-insoluble tau was boiled for 5 minutes at 95°C in 1X NuPage LDS Sample Buffer (ThermoFisher) + 4% BME. Samples were run on 4-20% Novex Tris-Glycine gels (ThermoFisher Scientific) for 50 minutes at 200 V and transferred onto PVDF membranes using the Trans-Blot Turbo Transfer system (BioRad). Membranes were blocked in TBS + 0.1% Tween-20 + 5% milk for 1 hour and incubated overnight at 4°C in primary antibody or for 2 hours at RT. Following washes in TBS + 0.1% Tween-20, blot was incubated in anti-mouse StarbrightBlue 700 (BioRad #12004158, 1:2500), anti-rabbit DyLight 800-conjugated secondary antibody (Cell Signaling Technology, 5151, 1:10,0000), and/or anti-goat IgG HRP (ROCKLAND, eB270, 1:1000) for 45 minutes. Signal was visualized using near IR fluorescence detection or chemiluminescence (ChemiDoc MP, BioRad). The following primary antibodies were used for immunoblotting: Tau 12 (Sigma-Aldrich, MAB2241, 1:2000); BR134 (in-house, rabbit polyclonal, 1:4000); RD3 (Sigma-Aldrich, 05-803, 1:1000); Tau 4R (Cell Signaling Technology; 30328); anti-GFP (Proteintech, 50430-2-AP, 1:2000); anti-HA.11 (Biolegend, 901516l; 1:2000); anti-GAPDH (ThermoScientific, MA5-15738; 1:2000); anti-CypA (Bio-techne; AF3589;1:1000).

### Full-length, wild-type tau HEK293 seeding assay

96 Well Black Polystyrene plates (Corning, CLS3603) were coated with Poly-D-Lysine (Gibco, A3890401) for 30 minutes at 37°C, before washing the plate 3x with 100μL / well of PBS. Tau assemblies were incubated at 2X final concentration in OptiMEM Reduced Serum Media (Gibco, 31985062) in a 50μL / well reaction mix with a 1:50 dilution of Lipofectamine 2000 (Invitrogen, 11668019) for 20 minutes at RT. Cells were trypsinzised and counted using Countess II Automated Cell Counter (Invitrogen). Cells were diluted to 40,000 cells / mL in OptiMEM, and 50uL of cells was combined with 50uL of tau assembly mixture per well for a final concentration of 1:100 Lipofectamine 2000, 1X tau, and 20,000 cells / well. Cells were returned to the incubator and, after 1 hour, 100μL / well of DMEM + 10% FBS was added to each well. Seeding was allowed to occur for 72 hours before analysis.

### Immunocytochemistry

HEK293 cells were fixed and permeabilized in 100% ice-cold methanol. Cells were blocked with PBS + 2% BSA and incubated in primary antibody at RT for 2 hours, or 4°C overnight. Cells were washed 3x in PBS and incubated in secondary antibody or streptavidin: Alexa Fluor 568 goat anti-mouse IgG (H+L) Highly Cross-Adsorbed (Invitrogen, A-11031, 1:1000); Streptavidin Alexa Fluor 488 (Invitrogen, S11223, 1:1000); Alexa Fluor 647 goat anti-rabbit IgG (H+L) Highly Cross-Adsorbed (Invitrogen, A-21245).

The following primary antibodies were used for immunocytochemistry FITC-anti-HA (Roche,11988506001, 1:200, discontinued); biotin-anti-HA (Roche, 12158167001, 1:500); AT100 (pT212/pS214) (1:500, MN1060).

Nuclei were stained with PUREBLU Hoechst 33342 (BioRad, 1351304). Amyloid filaments were labelled with Amytracker 680 (Ebba Biotech, 1:500).

### Indirect ELISA

Cell lysate was immobilized on 96-well High binding ELISA plates (Corning, 3590) in 1X ELISA Coating Buffer overnight at 4°C. Plate was washed 4x with PBS + 0.05% Tween-20 and blocked for 2 hours at RT in 5% milk in PBS. Blocking buffer was removed and HT7 primary antibody (MN1000) was diluted in 5% milk/PBS to a final concentration of 0.1μg/mL. Primary antibody was incubated on the plate for 1 hour at RT, then washed 4x with PBS + 0.05% Tween-20. Plate was incubated with HRP-goat anti-mouse IgG (H+L) (Invitrogen, A28177, 1:1000) in 5% milk for 1 hour at RT, then washed 4x with PBS + 0.05% Tween-20. TMB substrate (Cell Signaling Technology, 7004) was added for 30 minutes for signal detection and quenched with 0.16 M H_2_SO_4_ before reading at 450nM on a Varioskan LUX microplate reader (Thermo Scientific).

### Microscopy

Confocal images were acquired on a SP8 Lightning Confocal Microscope (Leica) with a 63x lens. Images for % seeding analysis were acquired on a Ti2 High Content Microscope (Nikon) with a 10x lens and analysed using NISElements software.

### Statistical Analysis

Statistical analysis was performed using GraphPad Prism software. All the data are shown as the mean ± SD.

## Acknowledgements

We thank the tissue donors and their families for their generous gift. We thank Dr Annelies Quaegebeur and Dr Olaf Ansorge of the Cambridge Brain Bank and Oxford Brain Bank. We are grateful to Dr Michel Goedert, MRC Laboratory of Molecular Biology, for constructive suggestions. M.H. was supported by a Studentship funded by Aprinoia Therapeutics Limited. This study was supported by a Sir Henry Dale Fellowship to W.A.M. jointly funded by the Wellcome Trust and the Royal Society (grant 206248/Z/17/Z) and by the Lister Institute for Preventative Medicine. Further support was provided by the UK Dementia Research Institute (award number UK DRI-2010) through UK DRI Ltd, principally funded by the UK Medical Research Council.

## Notes

### Competing Interest Statement

The authors have declared no competing interest.

